# DREAM-Stellar: Parallel and space efficient exact local alignment

**DOI:** 10.1101/2025.09.22.677748

**Authors:** Evelin Aasna, Simon Gene Gottlieb, Marcel Ehrhardt, Knut Reinert

**Author notes:** Contributing authors.

## Abstract

**Background:** Searching large genomic data sets for local alignments poses a computational challenge. A particular obstacle is the handling of repetitive sequences that appear in various contexts and incur a high runtime cost. For practical homology search, it is important to develop a specific but sensitive filter. Good filters reduce the search space before alignment without missing significant matches.

**Results:** We introduce DREAM-Stellar, a parallelized, updated version of the pairwise local aligner Stellar. The new aligner, DREAM-Stellar, is composed of four steps: preprocessing the queries and references, building a data structure for distributing the queries, computing in parallel the results and finally combining them. For distributing the queries we use the IBF data structure and a new prefilter for local alignments. We present our comparison of five local aligners on simulated and real genomic data and conclude that heuristic tools like BLAST miss a large percentage of significant local alignments or “drown” them in millions of less significant matches. The new version of Stellar is up to **900** times faster than its predecessor and can find all alignments between a pair of genomes in minutes. With that, the runtime of DREAM-Stellar is on par with tools like BLAST etc.

**Conclusions:** DREAM-Stellar is very practical and fast on very long sequences which makes it a suitable new tool for finding local alignments between genomic sequences under the edit distance model. The software is freely available for Linux and Mac OS X at https://github.com/seqan/dream-stellar

## 1 Background

For some basic problems in bioinformatics research there are suitable solutions that however warrant revisiting because of the constant increase in data volume to be analyzed. One such problem is the computation of good local alignments. The level of evolutionary conservation varies greatly along a genome. Local alignments between related sequences can point to conserved regions that are under selective pressure due to their fundamental role [1]. Local sequence alignments have been studied for over four decades, as evidenced by references like [2, 3]. Despite this extensive research, the problem continues to captivate the field.

For one, local alignments are employed to identify homologous regions in relatively short protein or nucleic acid sequences. This approach is used to find clipped alignments for divergent reads. There is a wealth of alignment tools such as Lambda3 [4], DIAMOND [5] and Minimap2 [6] that support short read input.

For local alignments in larger sequences, such as reads from third-generation sequencing or chromosomes of assembled genomes, searching requires careful reconsideration of the query prefiltering and parallelization. Some tools find the best alignment between whole genomes [7] [6]. However, genomic-scale alignments are only collinear for very closely related genomes, such as individuals of the same species. A genome-wide comparison of more distantly related samples necessitates the computation of local similarities to capture various complex rearrangements, e.g, translocation or duplication.

In this use case, instead of identifying specific homologous regions, the objective is to display all significant similarities between sets of sequences. Numerous tools have been developed for computing the set of pairwise local alignments. Among early examples, BLAT [8] prioritizes speed, while BWT-SW [9] emphasizes sensitivity and can report all local alignments. However, BLAT and BWT-SW are designed for queries of up to 200kb and 1Mb, respectively, and are unsuitable for searching between pairs of chromosomes. Efficient heuristics, particularly the BLAST family [10–12], are suitable for large-scale analyses but may miss significant alignments. Of the recent genome-wide local aligners, the single-threaded LASTZ [13] and multi-threaded LAST [14] use gapped seeds for increased sensitivity compared to the ungapped seeds in BLAST.

Stellar [15] addressed the genome-wide local alignment problem using a lossless ungapped k-mer filter to find all pairwise alignments. While at the time of publication, Stellar outperformed many methods, it most notably was not multithreaded and for some computations it took hours. In this paper, we propose DREAM-Stellar, which is a new version of Stellar. In DREAM-Stellar, we have reworked Stellar to add a distribution and prefiltering strategy (Fig. 1) and use the Interleaved Bloom Filter (IBF [16]) to index the reference database. DREAM-Stellar supports gapped seeds and applies a probabilistic k-mer filter to reduce the search space before computing alignments concurrently.

**Fig. 1.**
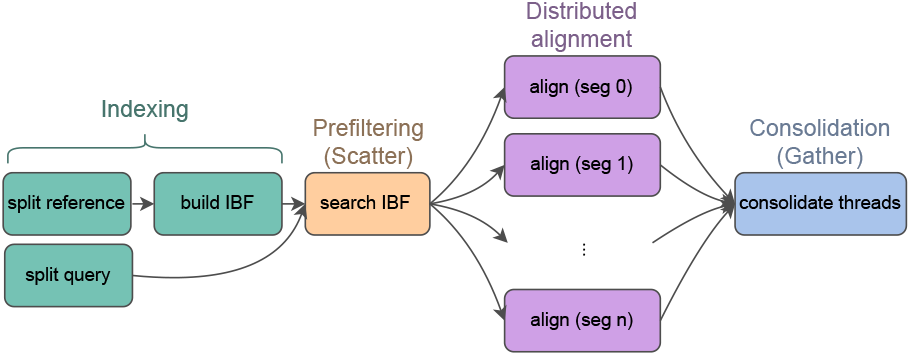
The input sequences are split into partially overlapping segments. Reference segments are processed to build an Interleaved Bloom Filter. Query segments are distributed and aligned in concurrent threads.

Finding all local alignments requires not only accuracy in terms of sensitivity, i.e., the development of precise tools that avoid overlooking regions of significant local similarity, but also computational efficiency when processing possibly repetitive long sequences. Between almost any set of sequences, there are many trivial local alignments. Due to this, it is necessary to apply a significance condition to define the alignments of interest. A maximal *E-value* threshold is a widely used criterion for choosing only significant local alignments. For each local alignment, the E-value is the expected number of alignments having an equal or better Smith-Waterman-like score that would appear in the input sequences by chance [11, 17]. The score underlying the E-value in LAST and LASTZ uses affine gap costs, whereas nucleotide BLAST by default uses linear gap costs that are suited for relatively low error rates. Stellar and DREAM-Stellar also output E-values based on linear gap costs, but do not apply an explicit E-value threshold.

Instead, DREAM-Stellar uses a maximal error rate *e*_*max*_, i.e, edit distance normalized by local alignment length as a score threshold and additionally requires a minimal alignment length *l*_*min*_. Alignments that score highly are also guaranteed to have a good E-value. A pairwise sequence alignment of length *n* containing *e* insertion, substitution or deletion columns is an *ε*(*l*_*min*_, *e*_*max*_)-match if *n* ≥ *l* and 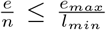. In DREAM-Stellar, we use a sensitive probabilistic k-mer filter to find *ε*-matches between large sequence sets. The k-mer size, which can have a profound effect on search speed and accuracy, we determine programmatically based on the input sequences and *ε*-match parameters.

## 2 Implementation

The Smith-Waterman algorithm [3] and the related Waterman-Eggert [18] algorithm are fundamental approaches to finding local alignments between two sequences that differ sufficiently from each other. However, the quadratic runtime of these algorithm severely restricts the size of sequences that can be aligned in a reasonable time. For this reason, it is common to preprocess reference sequences to build an index and to search the index for seed matches of the query before proceeding with alignment in a reduced search space. In DREAM-Stellar, we apply two rounds of increasingly specific prefiltering before using a banded version of the Waterman-Eggert algorithm to find local alignments between two sequence sets in reasonable time.

### Seeding

The widely adopted BLAST suite pinpoints regions of interest based on an exact word match. The LASTZ local aligner also relies on a word match but supports gapped words with some do-not-care positions. Gapped k-mers, represented as strings of 0’s and 1’s where zeroes stand for the gaps, are more sensitive than gapless seeds of the same length because gaps can accommodate edits. Also, the gapped seed 11011 is more specific than the longest gapless seed 11, which is guaranteed to have at least the same sensitivity. LAST also uses gapped seeds by default and adapts seed length if a word is found to be too frequent.

Like LASTZ and LAST, DREAM-Stellar supports gapped and ungapped words; however, seed matches in DREAM-Stellar are regions of multiple word hits within a region of the query sequence. Based on a chosen minimum length and maximum edit distance, DREAM-Stellar applies a threshold for the number of word hits in a query region. Because choosing k-mer size and threshold is crucial, we developed a probabilistic model to determine a suitable k-mer size and threshold for a particular dataset.

The k-mer shape can also be freely chosen, but by default, we use ungapped extensions of the seed 111010101101010111, which is a symmetrical derivative of the seed 111010010100110111 that was shown to be most sensitive by PatternHunter [19]. The asymmetrical PatternHunter seed is also used by LAST and LASTZ. Like Pattern-Hunter, we choose a gapped shape where adjacent k-mers share only 5 positions. Such seeds are sensitive because errors that fall into any position other than the 5 overlapping ones do not destroy both adjacent k-mers. Unlike PatternHunter, we consider the canonical k-mer, which at each seed position is the k-mer or its reverse complement, whichever has the smallest hash value. Because it is unclear if the forward or reverse complement k-mer will be chosen for a given position, we choose symmetrical seeds so that gaps are in the same position on both strands. Our chosen symmetrical seed guarantees that any adjacent canonical k-mers share only 5 positions (Table 1), unlike the PatternHunter seed, for which a reverse complement k-mer will share 8 positions with the next forward strand k-mer (Table 2).

**Table 1.**
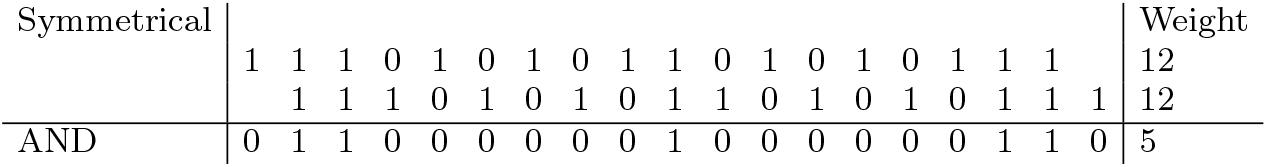
Shared positions for adjacent symmetrical seeds. The number of 1’s in a gapped seed is the k-mer weight. We find the number of positions shared by adjacent k-mers by a bitwise AND.

**Table 1.** Time and memory requirements for indexing the mouse genome. BLAST indexing is the fastest and least memory-intensive. LASTZ and Stellar do not create a reusable index before a search. The LAST index is the largest and slowest.

| Indexing (mouse) | Time | Memory | Size |
| --- | --- | --- | --- |
| BLAST | 23s | 360MB | 650Mb |
| LAST | 4min41s | 14.6GB | 13.5GB |
| LASTZ |  | n/a |  |
| Stellar |  | n/a |  |
| DREAM-Stellar<br>(gapped 19-mer) | 27s | 4.9GB | 4.3GB |

**Table 2.**
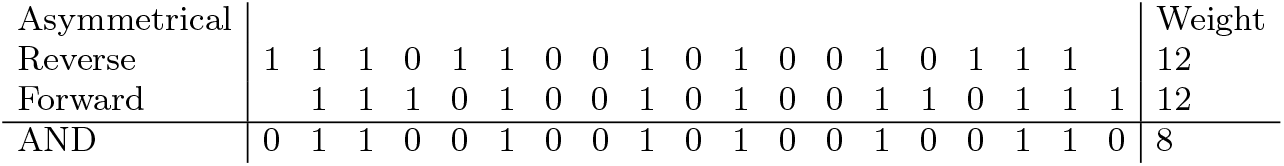
The number of shared positions between adjacent seeds from the reverse and forward strands for the shape 111010010100110111 that was shown to be the most sensitive weight 11 seed by PatternHunter. When using canonical k-mers, this asymmetrical seed has an overlap of up to 8 positions with its adjacent seed.

**Table 2.** Time and memory requirements for indexing the human genome.

| Indexing (human) | Time | Memory | Size |
| --- | --- | --- | --- |
| BLAST | 23s | 455MB | 766MB |
| LAST | 5min42s | 16.5GB | 16.5GB |
| LASTZ |  | n/a |  |
| Stellar |  | n/a |  |
| DREAM-Stellar<br>(gapped 23-mer) | 36s | 7.4GB | 6.5GB |

### Distribution strategy

In DREAM-Stellar, we split the reference and query sequences into segments of approximately equal length. The segments must partially overlap to find alignments that span the borders of the segments. The query segments are processed on concurrent threads. Initially, query segments are read in chunks of 10 million and scattered across the concurrent threads in “producer jobs”. All producer threads access the same reference index (IBF) to look up query k-mers in the initial prefiltering. Producer jobs associate each query segment with a set that contains zero or more reference segments.

The prefiltering results are used to create batches (carts) of segments matching the same reference region. Carts that reach the user-specified capacity are placed in the queue of full carts. Consumer threads dequeue full carts in a first-in, first-out manner, and create “consumer jobs”. Each consumer job builds a separate k-mer index over the batch of query segments. These full-text k-mer indices are used to reduce the search space before alignment further. For the alignment step, all consumer threads operate with the same copy of any reference or query sequence. Finally, the alignment results of all consumer jobs are gathered and consolidated to eliminate duplicates. A schematic of this approach is given in Fig. 2.

**Fig. 2.**
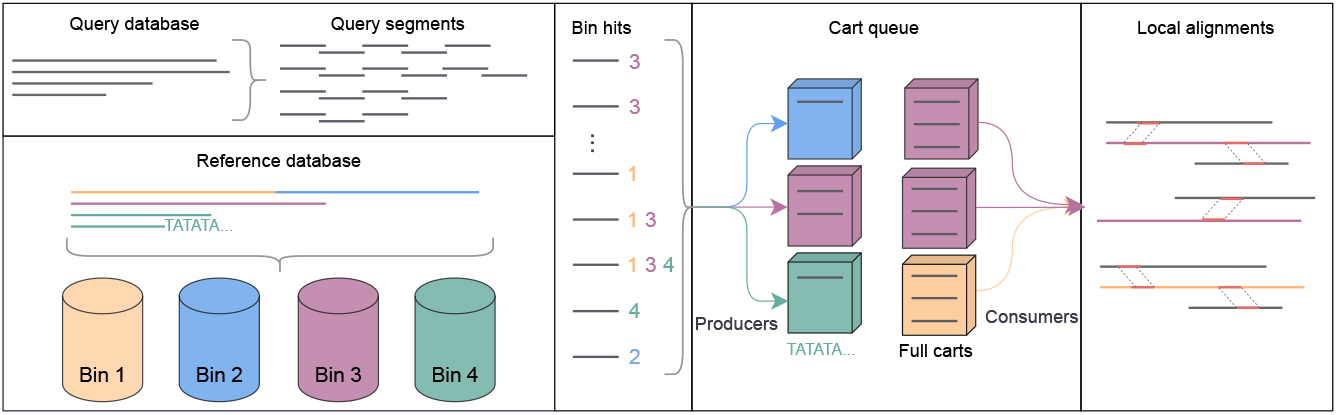
Pictured is a cart queue for an IBF of 4 bins. The maximum queue size as well as cart capacity are 3. The query with many bin hits is not mapped to the repetitive Bin 4.

Producing and consuming overlap in time. The first consumer job is launched when the first full cart gets added to the queue. In addition to cart capacity, the queue is parameterized by the maximum number of full carts in the queue. The memory footprint of the search can be reduced by choosing smaller parameter values. The cart capacity limits the size of any consumer job and the maximum number of filled carts limits the number of concurrent jobs.

### Prefiltering

Local alignments could start at any position in the reference and query sequences. To avoid an exhaustive alignment of the sequences, we apply two levels of increasingly specific prefiltering that result in candidate regions of local similarity. Splitting long sequences into partially overlapping segments allows processing linear sequencing data in parallel threads.

First, we reduce the search space for each query segment by discarding all reference regions that do not contain an *ε*-match. We omit the (*l*_*min*_, *e*_*max*_) notation for brevity. For this, we index a representative set of reference k-mers into an Interleaved Bloom filter (IBF) [20]. A membership lookup in the IBF outputs, with some false positive probability (FPR), the reference segments that contain a query k-mer. An upper limit for the FPR of an IBF k-mer query is set at index construction. Unlike a conventional seed-and-extend approach [11] [13] [14] that extends a single k-mer match, in DREAM-Stellar, seeds are regions of the query that contain more than some threshold number of k-mer hits. The threshold for the minimum number of shared k-mers that constitute a likely local match is calculated by a probabilistic model that extends the well-known k-mer counting lemma.

IBF prefiltering is an efficient way to discard reference regions that do not match some query segment; however, not all IBF hits constitute an *ε*-match. Next, the set of IBF hits for each reference segment is further verified in the more specific and more costly SWIFT [21] filter. For this, the query segments are indexed using a k-mer index that yields the location of all query k-mers. For a detailed description of the lossless SWIFT filter, refer to the original Stellar paper [15]. In DREAM-Stellar, we split the input sequences into segments and reduce the search space using the IBF before the distributed execution of SWIFT filtering and Stellar alignment.

### Parameter deduction

Generally, shared k-mers suggest sequence similarity. In IBF prefiltering, we count the shared k-mers between query regions and reference bins, and use the shared k-mer count to gauge whether the reference contains an *ε*-match for the query. Sequence pairs that share no or only a few k-mers are discarded from further search. This is important for achieving a low running time.

The well-known k-mer counting lemma gives a lower bound for the number of k-mers shared by the query and reference regions if an *ε*(*l, e*)-match exists. The k-mer counting lemma states that given two sequences that share a local alignment of length *l* that differ by *e* edit operations, the two sequences will share at least *l* − *k*(*e* + 1) −1 k-mers. Intuitively, when the edit distance increases, the shared k-mer count for a fixed *k* decreases and eventually becomes zero. For a high edit distance, *k* must be small to yield a positive threshold for filtering.

Conversely, when k is small and a sequence is sufficiently long, the sequence likely contains every possible subsequence of length k. The left panel of Fig. 3 shows that both the real and randomly sampled 143Mb genomes contain every canonical 11-mer. Ideally, prefiltering could attribute query k-mers only to the position of an *ε*-match in the reference. However, k-mers that stem from repeat regions appear many times across the genome. Moreover, for a small k, most k-mers appear many times in the genome. This leads to false positive IBF hits, which are k-mer-based matches that do not correspond to an *ε*-match.

**Fig. 3.**
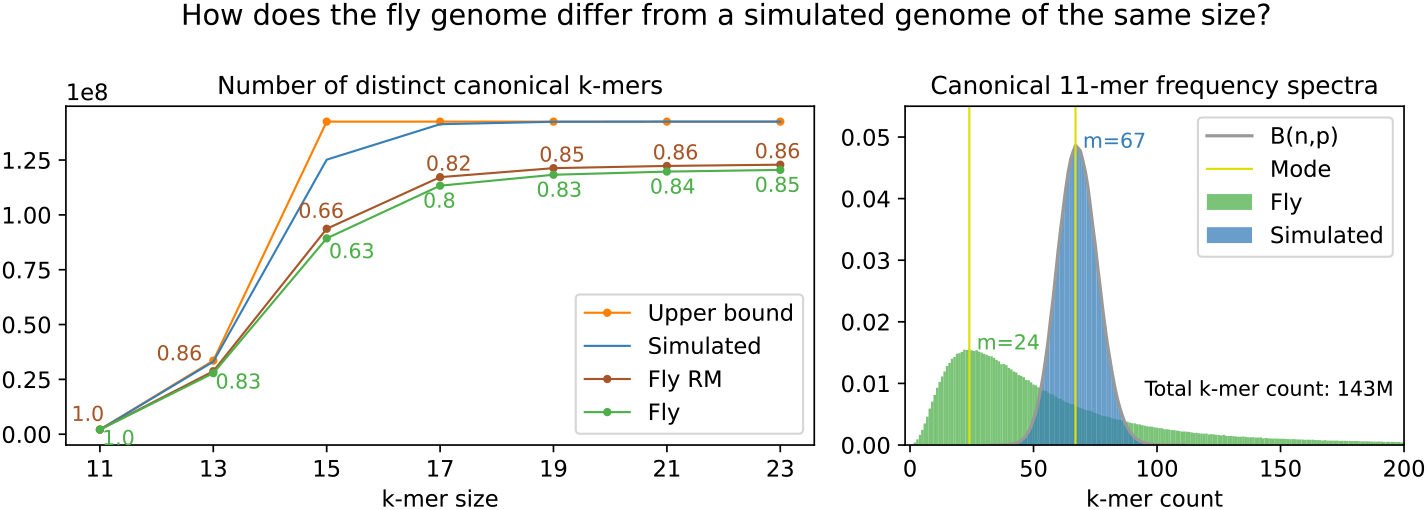
Genomes contain repetitive elements that are not represented by a simple sequence simulation model. **Distinct count:** The fruit fly (Release 6) reference genome contains fewer distinct k-mers compared to a simulated genome of equal size, where each of the 143 million DNA bases is sampled independently from a uniform distribution. The effective size of the fly genome is approximately equal to sequence length for 11-mers and varies between 63% and 85% for longer k-mers. Repeat masking (RM) introduces random sequence in place of low-complexity sections and slightly increases the effective sequence size. **Frequency spectra:** Although almost all 11-mers appear at least once in both the simulated and real fly genomes (effective size coeff 1.0), the 11-mer abundance distribution is very different. The long right tail of the fly distribution is not pictured; the most frequent 11-mer, the A homopolymer, appears 89k times.

To estimate the suitability of different k-mer sizes, we model a sequence of length *L* as the set of k-mers. A sequence of length *L* has a total of *L* − *k* + 1 k-mers. Let *X* be the random variable representing the number of occurrences of some canonical k-mer when sampling a uniform distribution *L* − *k* + 1 times. There are |*σ*| ^*k*^ distinct k-mers in an alphabet of size |*σ*| ; half of these can be chosen as the canonical k-mer. Thus, *X* has a binomial distribution and a success probability of 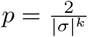 (right panel Fig. 3). Strictly, only bases and not k-mers are sampled independently in sequence simulation, but the dependence on adjacent overlapping k-mers is negligible when *k* ≪ *n*.

The expected number of occurrences for a canonical k-mer in a random DNA sequence is 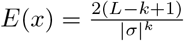. We estimate the probability of a k-mer match in the case of independent and random sequences, considering the size of the reference database as *P* (*x*) = *min*(1, *E*(*x*)).

For a simulated fly genome (143Mb), we expect a 11-mer to appear 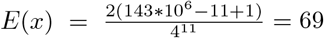 times. We estimate the probability of a spurious 11-mer match to be *P* = *min*(1, 69) = 1 and call this the false positive probability of a k-mer. Because any 11-mer is expected to occur in the 143Mb sequence just by chance, 11-mers are not suitable to reduce the search space in prefiltering. Refer to the appendix for further details on the combinatorial approach to extend the spurious k-mer match probability to find the probability of at least the threshold number of k-mers matching spuriously.

For the extreme case of a large reference sequence and a high error allowance, it might not be possible to find a k-mer size that does not appear by chance but also yields a positive k-mer lemma threshold. Luckily, the k-mer lemma threshold is the lower bound for the number of shared k-mers in case errors are distributed uniformly across the alignment.

The number of ways in which errors can be distributed across the local alignment is the number of combinations 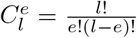. Assuming that all error configurations are equally likely, the probability of encountering the uniform error configuration becomes negligibly small as the number of edits and the epsilon length increase. Further, because edits are likely to appear in clusters, i.e., indels are common, any sequences sharing a high edit distance *ε*-match most likely share more than the k-mer lemma threshold number of matches.

Given an *ε*-match definition, we estimate the false negative rate of some k-mer size and threshold by finding the fraction of error configurations that preserve fewer than the threshold shared k-mers. For this, we use a dynamic programming (DP) algorithm to precalculate the number of false negative error configurations for a wide range of parameter values once (ca 1sec and 6MB) and reuse the DP table when finding the best k-mer size and threshold for any dataset. Refer to the appendix for details on the 3D dynamic programming algorithm.

To deduce suitable parameters for some dataset size and *ε*-match definition, we minimize the sum *f* (*k, t*) = *FPR* + *FNR*.

### Effective sequence size

The distinct k-mer count *C* in a sequence of length *L* is at most 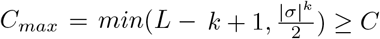. Except for the short k-mer case where 4^11^ ≪ 140 ∗ 10^9^ − 11 + 1 and all 11-mers appear by chance, there are fewer distinct k-mers in the fruit fly reference genome compared to a randomly simulated genome of equal length (Fig. 3) *C*_*max*_ *> C*. The false positive rate estimate in our threshold model depends on the probability of drawing a random element (k-mer) from the reference sequence.

Because a real genome contains fewer distinct k-mers than is expected for a sequence of that size, we introduce the measure of effective sequence size 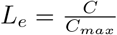. The effective sequence size of real genomes is less than expected from an independent and random sequence model.

The left panel of Fig. 3 shows that for sequences of size 143Mb, the number of distinct k-mers grows with k-mer sizes and plateaus around *k* = 19, meaning that there is little improvement in specificity when searching 19 or 23-mers in the fly genome. Because smaller k-mers are more sensitive when searching with errors, 19-mers are preferred in this case.

However, the k-mer spectra for reference genomes are not known. Counting the number of distinct k-mers for each reference dataset is not feasible and we correct the effective distinct k-mer set by a constant factor *s*_*e*_ = 0.65, which increases the false positive probability of a k-mer.

### Repeat filtering

It is well known that genomes contain repeats. There are genomic motifs such as tandem repeats, SINES, LINES etc. that are abundant across a given genome. Computing local alignments across repeats is usually not done, because it is of little interest and lets the run time explode. Hence, tools such as RepeatMasker [22] or Dust [23], which is the default in BLASTn, are used to exclude such regions.

In DREAM-Stellar, we avoid verifying all redundant repetitive regions by favoring highly variable regions. If a query has many IBF matches across the reference, then we preferentially search the less repetitive segments. We quantify the repetitiveness of a sequence segment with a simple measure. Given a segment of length *L*_*i*_ and the number of distinct k-mers in the sequence *C*_*i*_, the variability *V*_*i*_ of a segment is the fraction of distinct k-mers among all k-mers, 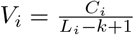.

Fig. 4 shows *R*_*i*_ which is the repetitiveness of segments across the human reference genome, *R*_*i*_ = 1 − *V*_*i*_. Repetitiveness varies from 0, for segments where all 19-mers are unique, to 1, for segments that only contain a single 19-mer, which is mostly the case for stretches of unknown N bases. When a query segment is abundant and matches more than 50% of reference segments, we favor variable regions of the reference and avoid an exhaustive search. Queries that match only a few repetitive reference regions are searched in all of them.

**Fig. 4.**
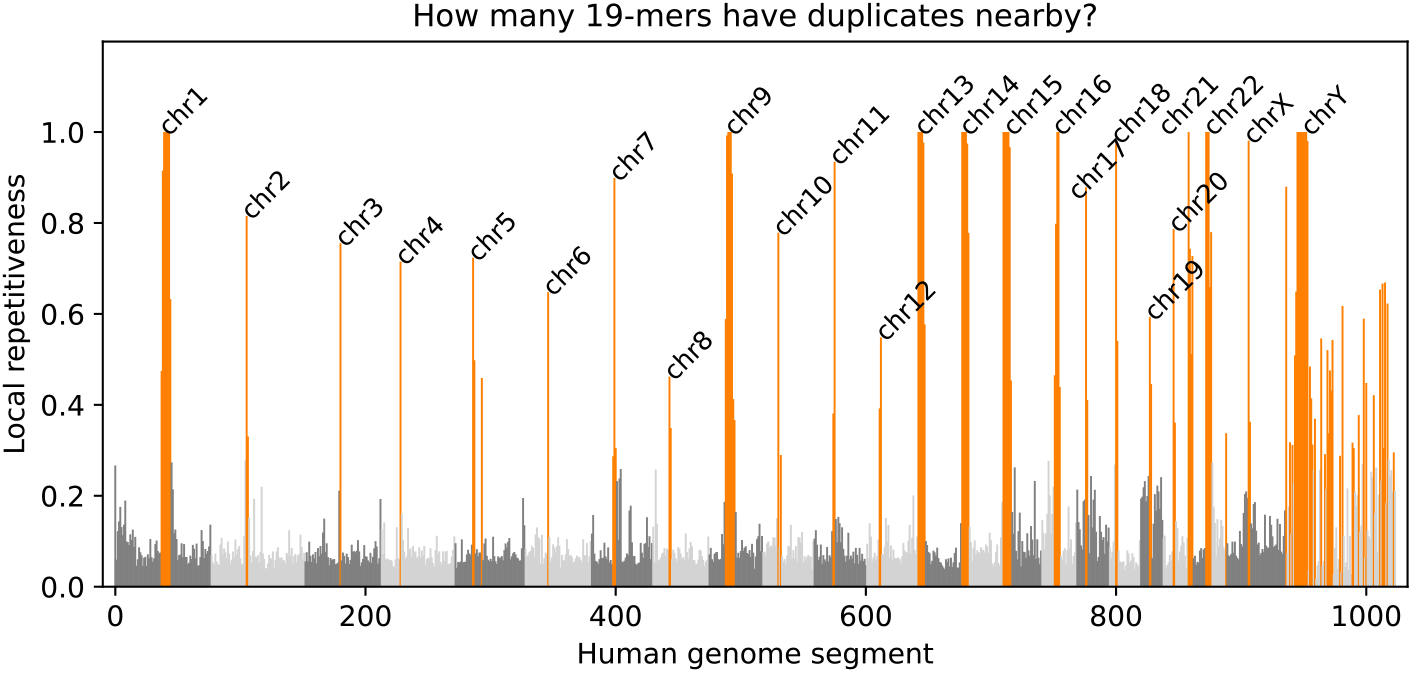
We divided the human reference genome into 1024 segments and found the segment local repetitiveness of 19-mers. The main chromosomes are colored in alternating gray tones. Alternative scaffolds of the reference genome are pictured downstream of chrY in light gray. The top 25% of highly repetitive regions are shown in orange. Many highly repetitive segments are in the centromere of the corresponding chromosome. The Y chromosome is also particularly repetitive.

## 3 Results

We compare DREAM-Stellar against state of the art local alignment programs, namely nucleotide BLAST[10] (version 2.12.0), LASTZ[13] (version 1.04.22) and LAST[14] (version 1595) in terms of speed and sensitivity and show that some of the tools are prohibitively slow or miss many significant local alignments that can be detected with DREAM-Stellar.

Benchmarks were run on a server with a dual-socket AMD EPYC 9454 48-Core processor system and 1007GB of available RAM. Time is wall clock time. Where applicable, the search is run concurrently on 32 threads; LASTZ and Stellar do not support parallel execution. To quantify the effect of prefiltering before alignment, we compare DREAM-Stellar to a naively parallelized version of Stellar, where reference sections are searched for matches for all queries in parallel.

### Simulated data

We first assess the default behavior of the tools on simulated data where the true set of local alignments is known. For this, we simulated pairs of random sequences with lengths of 50MB and 250MB, approximately equal in length to human chr21 and chr1, respectively.

We then sampled one sequence in the simulated chr21 pair for 26k (or 131k for simulated chr1) local alignments varying in length between 50 and 250bp and containing 0-10% of errors distributed uniformly randomly across the alignment. Finally, we inserted the simulated local alignments into the other simulated chromosome in the pair at random positions without overlapping. On average, there is one local alignment for each 2000bp of sequence.

Table 3 and Table 4 show the time and accuracy when searching between the simulated pairs of sequences for local alignments. We compare BLAST with the nucleotide default parameter values of a 28bp word size, with the k-mer lemma derived minimum word size guaranteed to find all local matches of length at least 50bp. These word sizes are 49, 25, 16, 12, 10, and 8 for 0-5 errors.

**Table 3.** Local alignments between two simulated sequences of length 50Mb. With the default word size of 28, BLAST misses up to 47% of highly significant local alignments. For space reasons, the RAM peak is only shown for two error rates.

| Error rate | 0% |  | 2% |  |  | 4% |  |
| --- | --- | --- | --- | --- | --- | --- | --- |
|  | Time | Missed | Time | RAM | Missed | Time | Missed |
| BLAST (default) | 9s | 0% | 9s | 364Mb | 0% | 9s | 3% |
| BLAST (exact) | 8s | 0% | 9s | 366Mb | 0% | 16s | 0% |
| LAST | 2m54ss | 0% | 2m53s | 546Mb | 0% | 2m54s | 0% |
| LASTZ | 28m56s | 0% | 29m2s | 561Mb | 0% | 28m50s | 0% |
| Stellar (single) | 1m10s | 0% | 1m21s | 3.2Gb | 0% | 13s | 0% |
| Stellar (dist) | 9s | 0% | 10s | 100Gb | 0% | 9s | 0% |
| DREAM-Stellar | 7s | 0% | 7s | 544Mb | 0% | 4s | 0% |

| Error rate | 6% |  | 8% |  |  | 10% |  |
| --- | --- | --- | --- | --- | --- | --- | --- |
|  | Time | Missed | Time | RAM | Missed | Time | Missed |
| BLAST (default) | 9s | 13% | 9s | 369Mb | 29% | 9s | 47% |
| BLAST (exact) | 3m8s | 0% | 13m42s | 416Mb | 0% | 2h13m | 0% |
| LAST | 2m53s | 0% | 2m53s | 544Mb | 0% | 2m53s | 0% |
| LASTZ | 28m43s | 0% | 29m2s | 557Mb | 0% | 28m48s | 0% |
| Stellar (single) | 2min36s | 0% | 7h27min | 1.4Gb | 0% | 10h36m | 0% |
| Stellar (dist) | 12s | 0% | 27m30s | 38Gb | 0% | 36m12s | 0% |
| DREAM-Stellar | 4s | 0% | 4s | 550Mb | 0% | 6s | 0% |

**Table 4.** Local alignments between two simulated sequences of length 250Mb. With the default word size of 28, BLAST misses up to 48% of highly significant local alignments. LAST higher runtime for 6% error rate was consistent through multiple repetitions and accounting for I/O effects. Due to being much slower than default BLAST, LAST and DREAM-Stellar, LASTZ was only executed once. For high error rates, Stellar and exact BLAST search were not finished after 12h. For space reasons, the RAM usage peak is shown only for two error rates.

| Error rate | 0% |  | 2% |  |  | 4% |  |
| --- | --- | --- | --- | --- | --- | --- | --- |
|  | Time | Missed | Time | RAM | Missed | Time | Missed |
| BLAST (default) | 1min21s | 0% | 1min20s | 1.5G | 0% | 1min20s | 4% |
| BLAST (exact) | 56s | 0% | 1min30s | 1.5G | 0% | 4min11s | 0% |
| LAST | 15min28s | 0% | 15min34s | 1.7G | 0% | 15min42s | 0% |
| LASTZ | 11h | 0% | 11h | 2.2G | 0% | 11h | 0% |
| Stellar (single) | 3h37min | 0% | 46min | 14G | 0% | 3min30s | 0% |
| Stellar (dist) | 8min59s | 0% | 2min41s | 430G | 0% | 46s | 0% |
| DREAM-Stellar | 7min20s | 0% | 2min30s | 2.5G | 0% | 40s | 0% |

| Error rate | 6% |  | 8% |  |  | 10% |  |
| --- | --- | --- | --- | --- | --- | --- | --- |
|  | Time | Missed | Time | RAM | Missed | Time | Missed |
| BLAST (default) | 1min20s | 13% | 1min19s | 1.5G | 30% | 1min17s | 48% |
| BLAST (exact) | 1h16min | 0% | 5h30min | 1.6G | 0% | >12h | n/a |
| LAST | 20min6s | 0% | 18min40s | 1.7G | 0% | 18min3s | 0% |
| LASTZ | 11h | 0% | 11h | 2.2G | 0% | 11h | 0% |
| Stellar (single) | 3h57min | 0% | >12h | n/a | n/a | >12h | n/a |
| Stellar (dist) | 10min47s | 0% | 6h4min | 168G | 0% | 6h42min | 0% |
| DREAM-Stellar | 39s | 0% | 63s | 2.6G | 0% | 4min24s | 0% |

Our assessment on simulated data found that BLAST with the default parameters misses up to 47% of highly significant alignments. We required at least 10bp overlap to count an alignment as found. By default, nucleotide BLAST searches for exact 28bp seeds, which are not sensitive to alignments with a uniform error configuration. Conversely, all significant alignments contain short seed matches, but these short words may be ubiquitous and unable to reduce the search space. This is why BLAST runtime significantly increases for shorter word sizes. The original Stellar suffers from a similar effect at higher error rates because it also relies on exact word matches to direct the alignment using the SWIFT filter. LASTZ and LAST both use gapped seeds and have full sensitivity on simulated data, but LASTZ is slow because queries are not processed concurrently. DREAM-Stellar is faster than the other tools. Especially for higher error rates, there is a notable speed up compared to the single threaded original Stellar and the naively distributed version that does not use IBF filtering. Similarly, a naively parallelized version of Stellar uses two orders of magnitude more RAM than DREAM-Stellar.

In the tested edit distance range, DREAM-Stellar takes longer to find exact matches than approximate matches (Table 4). Exact search is slow, because the time complexity of part of the X-drop alignment algorithm in Stellar depends on 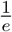, which is a large number for *e* = 0. Because most biological applications require approximate alignments, we have not implemented a separate subroutine for exact string matching.

The best parameters for the k-mer filter are calculated in the DREAM-Stellar workflow. For a 50MB dataset and *ε*(50, 5)-matches the cost minimum across the parameter space is at *Cost*(12, 5) = 0.27. The cost is the sum of *FNR*(12, 5) = 0.20 and *FPR*(12, 5) = 0.07, these estimates are based on ungapped k-mers. We searched both simulated references for at least 5 k-mer matches for the gapped shape 1110100101001101 (weight 12 Fig. 5) and found all matches. Even for the worst case, i.e., a simulated dataset where edits are distributed uniformly randomly, the FNR estimate is a loose upper bound for the actual number of missed matches. Symmetrical gapped k-mers are more sensitive than ungapped k-mers of the same weight, but a tighter bound for the sensitivity of gapped k-mers is an open question.

**Fig. 5.**
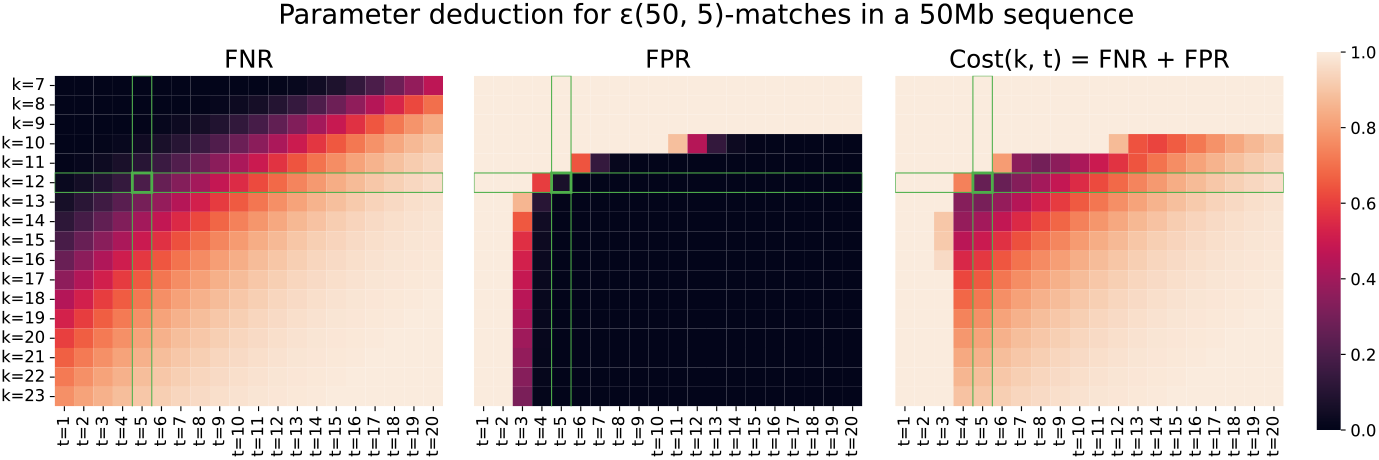
We deduce the parameters for searching a 50Mb reference sequence for *ε*(50, 5)-matches by minimising the sum *Cost*(*k, t*) = *FNR* + *FPR* over the k-mer weight *k* ∈ [7, 23] and shared k-mer count threshold *t* ∈ [1, 25]. Values close to 0 are better. The FNR estimate increases linearly with the k-mer weight and threshold. The FPR decreases exponentially with an increase in k-mer weight or threshold. The cost is minimal for 12-weight k-mers and a threshold of 5 *Cost*(12, 5) = 0.27.

### Reference genomes

Next, we present a comparison of the genome-wide local alignment between two pairs of reference sequences. We assessed the tools on the human (GRCh38), house mouse (GRCm39) and fruit fly (Release 6) reference genomes. The human and mouse genomes are less diverged than human and fly and similar in size, namely 3.1 and 2.6GB, respectively. The fruit fly genome is highly diverged from mammals and is 143MB in size. Due to higher divergence, we defined a shorter 50bp minimum length in the mouse vs fly comparison compared to the 100bp min length in the human vs mouse comparison. To make results comparable between tools, we turned off native repeat masking and used the Dust repeat masker (version 5.5.6) [23] to mask low-variability regions of the query before local alignment. For the following genome-wide benchmarks, we masked low-complexity regions using random sequence. The left panel of Fig. 3 shows that repeat masking (RM) slightly increases the number of distinct k-mers in the resulting sequence. We used the set of alignments from the slow but provably exact original Stellar aligner as the truth set. Missed matches are *ε*-matches for which no alignment overlaps by at least half of the minimum length (25bp for 50bp min length). Additionally, we listed BLAST, LAST, and LASTZ matches that are less significant than *ε*-matches in terms of length and edit distance.

### Mouse vs fly

A default BLAST *word* = 28 search between the mouse and fly genomes yields 5k matches. These include 158 matches that are less than 50bp long and contain more than 2 errors (Fig. 6 B). Even so, BLAST misses 19% of highly significant *ε*(50, 2)-matches (Table 5). Both the short BLAST matches and missed *ε*-matches occur across multiple chromosomes. For a more sensitive search, we decreased the BLAST seed length (*word* = 24) and found all but 11% of *ε*-matches, but also introduced 487 less significant matches.

**Table 5.** All local alignments between the M. musculus (GRCm39) and D. melanogaster (Release 6) reference genomes. BLAST, LAST, Stellar (dist) and DREAM-Stellar were run on 32 threads; LASTZ and Stellar do not support multithreading. Missed matches are *ε*-matches of at least 50bp with up to 1, 2 or 3 errors for which there was no local alignment that overlaps by at least 25bp. Stellar and DREAM-Stellar filter matches based on the error rate; for the other tools, the error count is relevant only in the accuracy assessment. By default, LASTZ and LAST use the gapped seed 1110100110010101111 from PatternHunter. For this benchmark, we also used an extension of this non-symmetrical seed for DREAM-Stellar.

| Mouse vs fly | Time<br>(RAM) | Matches | Missed<br>1 error | Time<br>(RAM) | Matches | Missed<br>2 errors | Time<br>(RAM) | Matches | Missed<br>3 errors |
| --- | --- | --- | --- | --- | --- | --- | --- | --- | --- |
| BLAST<br>word size 28 | 44s<br>(1.7G) | 5078 | 0% | 44s | 5078 | 19% | 44s | 5078 | 29% |
| BLAST<br>word size 24 | 55s<br>(1.8G) | 9k | 0% | 55s | 9k | 11% | 55s | 9k | 18% |
| BLAST<br>word size 16 | 3m30s<br>(2.8G) | 24k | 0% | 3m30s | 24k | 0% | 3m30s | 24k | 0% |
| LAST<br>init matches 10 | 2m32s<br>(14.8G) | 44k | 30% | 2m32s | 44k | 23% | 2m32s | 44k | 21% |
| LAST<br>init matches 100 | 14m22s<br>(15.3G) | 68k | 0% | 14m22s | 68k | 0% | 14m22s | 68k | 0% |
| LASTZ<br>step length 1 |  |  |  | >24h |  |  |  |  |  |
| LASTZ<br>step length 3 | 2h55m<br>(10G) | 69k | 0% | 2h55m | 69k | 1% | 2h55m | 69k | 1% |
| LASTZ<br>step length 30 | 19m50s<br>(6.9G) | 19k | 2% | 19m50s | 19k | 9% | 19m50s | 19k | 12% |
| Stellar (single)<br>min length 50 | 5m48s<br>(9.7G) | 1638 |  | 16m51s<br>(9.7G) | 3551 |  | 14h45m<br>(5.8G) | 5179 |  |
| Stellar (dist)<br>min length 50 | 44s<br>(225G) | 1638 | 0% | 1m35s<br>(227G) | 3551 | 0% | 1h15m<br>(101G) | 5179 | 0% |
| DREAM-Stellar<br>min length 50<br>gapped 23-mer | 17s<br>(10.4G) | 1638 | 0% | 26s<br>(12.7G) | 3551 | 0% | 1m1s<br>(13.5G) | 5134 | 1% |

**Fig. 6.**
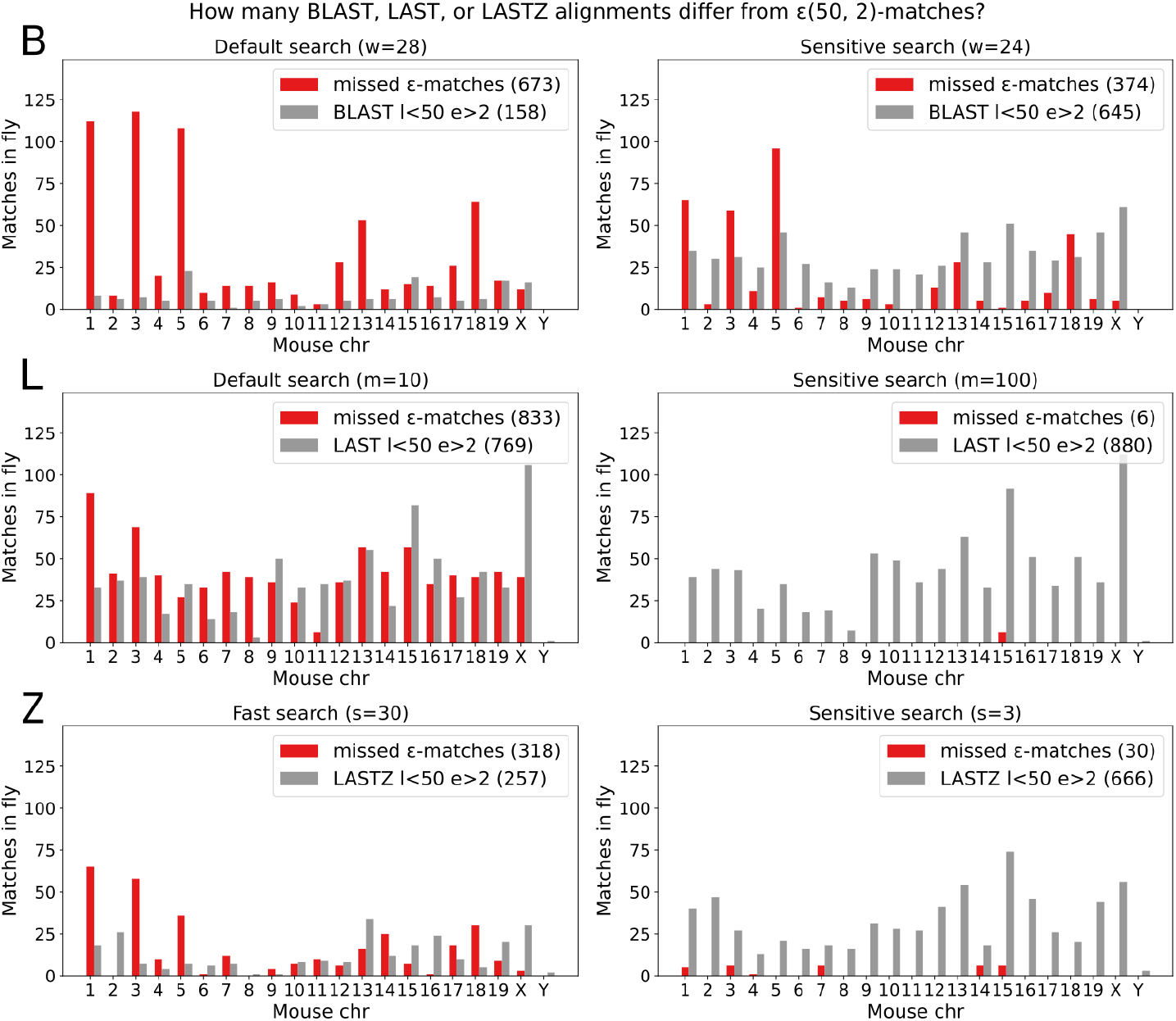
The tools in the comparison produce partially overlapping but distinct sets of alignments between mouse and fly. The number of alignments depends on some input parameters. By default, nucleotide **BLAST** initial matches are exact seeds of length *w* = 28. With the default word size, BLAST misses 673 *ε*(50, 2)-matches and finds 158 less significant matches that are less than 50bp long and contain more than 2 edits. BLAST sensitivity can be increased by reducing the word size. In **LAST**, there is a limit to the number of initial matches retained per query position. With the default initial match limit *m* = 10, LAST misses 833 *ε*(50, 2)-matches and finds 769 less significant matches that are less than 50bp long and contain more than 2 errors. LAST sensitivity can be increased by increasing the number of initial matches per query position. By default, **LASTZ** considers every query position in seeding, which is very slow. For a slightly sparser seeding of every 3rd position *s* = 3, LASTZ is still sensitive but misses 30 *ε*(50, 2)-matches and finds 666 less significant matches that are less than 50bp long and contain more than 2 errors. Search can be further sped up by increasing the seeding step. For all the considered parameters, runtime increases with sensitivity.

In this mouse vs fly comparison, BLAST has full sensitivity with *word* = 16bp. This is different in simulated data, where errors have a uniformly random distribution and 12bp and shorter seeds have full sensitivity for *ε*(50, 3) alignments.

A default LAST search between the mouse and fly genomes results in 44k matches. These include 769 matches that are less than 50bp long and contain more than 2 errors (Fig. 6 L). Even so, LAST misses 23% of highly significant *ε*(50, 2)-matches (Table 5). In sensitive mode, we increased the maximum number of initial matches per query position (m) and found all but 6 *ε*(50, 2)-matches and introduced 111 less significant matches.

By default, LASTZ considers every reference position as a seed. Because the default LASTZ with a seeding *step* = 1 was very slow in the simulated benchmark, we increased the seeding step parameter for the next benchmarks. There was little loss in sensitivity with *step* = 3, but LASTZ was still two orders of magnitude slower than the multithreaded tools (BLAST, LAST, DREAM-Stellar) in the comparison. The LASTZ *step* = 3 search between the mouse and fly reference genomes yielded 69k matches (Fig. 6 Z). These include 666 that are less than 50bp long and contain more than 2 errors (Table 5). As expected, increasing the seeding step further to *step* = 30 reduced the runtime but resulted in missing 9% of *ε*(50, 2)-matches.

DREAM-Stellar was again fastest and found all *ε*-matches for all but the highest error rate, where 1% of matches were missed due to applying a heuristic k-mer count threshold. Besides the naively parallelized Stellar that has a high memory footprint of *>* 100GB, all the other analyses require *<* 16GB and fit into the main memory of a modern personal computer.

Because of the differences in seeding, alignment scoring, and filtering between the tools, the resulting alignment sets differ. BLAST and LAST, in particular, miss ca 20% of highly significant *ε*-matches while producing hundreds of less significant alignments that are short and highly divergent.

### Human vs mouse

Because the mouse and human genomes are more closely related than the mouse and fly genomes, we increased the minimum match length to 100bp when searching for *ε*-matches with up to a 6% error rate. The maximum ungapped word size guaranteed to find all *ε*(100, 6)-matches is 16. BLAST *word* = 16 did not finish after 24h (see Table 6). With the default parameter values, BLAST and LAST were slower than all tested DREAM-Stellar instances. Unsurprisingly, the runtime of the single-threaded LASTZ is much slower compared to the tools run on 32 threads and LASTZ did not finish after 24h.

**Table 6.** All local alignments between the H. sapiens (GRCh38) and M. musculus (GRCm39) reference genomes. Due to the very high number of alignments found by BLAST and LAST, we did not compare these to the set of *ε*-matches. Stellar alignment did not finish due to memory restrictions. The DREAM-Stellar parameter set is shown for all error rates.

| Human vs mouse | Time | RAM | Matches | Additional parameters |
| --- | --- | --- | --- | --- |
| BLAST<br>word size 28 | 1h28m | 17G | 44M | E-value <10 |
| BLAST<br>word size 24 | 16h20m | 40G | 106M | E-value <1e-10 |
| BLAST<br>word size 16 | >24h |  |  | E-value <1e-10 |
| LAST<br>init matches 10 | 44min | 74G | 251M |  |
| LAST<br>init matches 100 | 5h20min | 393G | 1.7G |  |
| LASTZ<br>step length 3 | >24h |  |  |  |
| Stellar<br>1 error |  | OOM |  | min length 100 |
| DREAM-Stellar<br>1 error | 6m22s | 18G | 20k |  |
| DREAM-Stellar<br>2 errors | 6m25s | 18G | 34k |  |
| DREAM-Stellar<br>3 errors | 6m31s | 20G | 52k |  |
| DREAM-Stellar<br>4 errors | 7m36s | 32G | 76k | gapped 22-mer<br>11110110101101101111 |
| DREAM-Stellar<br>5 errors | 7m32s | 32G | 106k | min length 100<br>4096 bins |
| DREAM-Stellar<br>6 errors | 9m13s | 33G | 143k | 550k segments |

Table 6 shows a large discrepancy in the number of DREAM-Stellar *ε*-matches (143k) and the local alignments from LAST (251M) and BLAST (44M). Unlike BLAST, LASTZ, and LAST, Stellar defines the set of valid local alignments using a minimum length and maximum edit distance criteria. Although all DREAM-Stellar *ε*(100, 6)-matches in the human and mouse comparison have a low E-value of *<* 10^−28^, an E-value cutoff yields a partially overlapping but distinct set of alignments compared to any *ε*-match definition. The *ε*(100, 6)-matches yield a set of alignments that form contiguous blocks between human and mouse chr1 (Fig. 7). In contrast to the maximum normalized edit distance of 6% in DREAM-Stellar, the highest edit distance in this region is 32% among BLAST matches and 47% among LAST matches. As a result of a liberal similarity criteria, the default BLAST and LAST output alignment files are 3.1GB and 20GB in size, respectively.

**Fig. 7.**
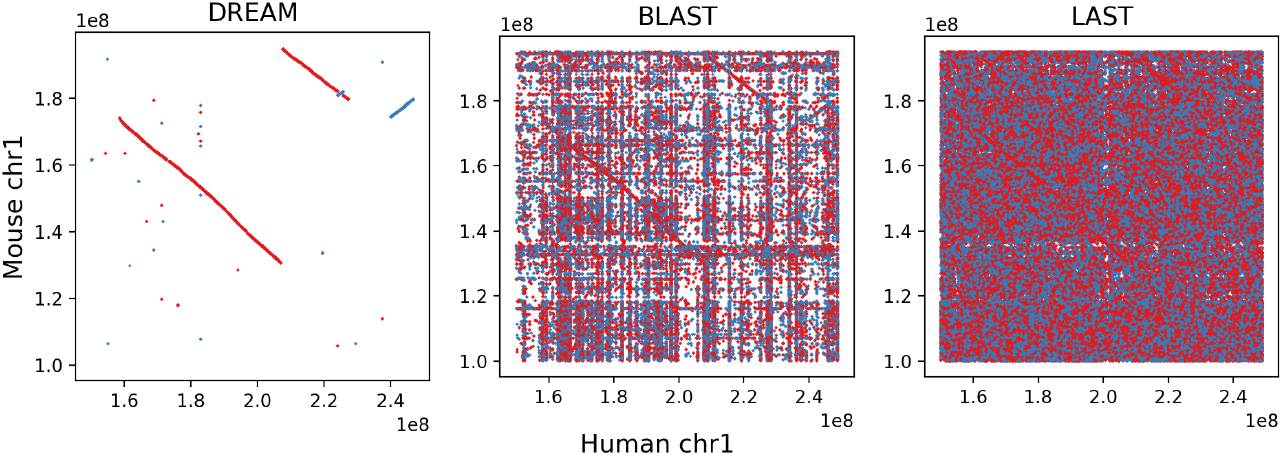
Local alignments between sections of 1p (short arm of chromosome 1) in mouse and human. Alignments between the forward strands are in blue and inverted mouse alignments are in red. The set of DREAM-Stellar *ε*(100, 6)-matches illustrates the large contiguous blocks of homology between chromosome 1 in mouse and human. BLAST (*w* = 28 and *E <* 10) yields many matches, with some regions appearing often across the homologous chromosome. LAST (*m* = 10) aligns most regions of human chr1 to most regions of mouse chr1.

For both genome-wide comparisons, the number of DREAM-Stellar matches incrementally increases with the error allowance. Despite the shorter minimum length of 50bp, the mouse and fly comparison (Table 5) resulted in fewer short local alignments than the search between the pair of mammalian genomes (Table 6). DREAM-Stellar is faster than the other tested tools and has a memory footprint comparable to LAST and BLAST. Compared to LAST and BLAST, DREAM-Stellar yields fewer, but highly significant alignments that fit the *ε*-match criteria.

### Structural variant detection

Finally, we used DREAM-Stellar in a structural variant (SV) detection workflow and compared the local alignments we found with previously published results [24] [25]. SVs are large abberations of ≥50bp that can complicate read mapping.

The human structural variant project(HSVP) reported 108k variants with a mean length of 1061bp (std dev 54990, median 164bp). We reanalyzed PacBio HiFi sequencing data for 4 samples (from 2 men and 2 women HG00731, HG00732, NA19238, NA19239) from Phase II of the Human Structural Variation Project [26]. First, we map the HIFI reads using pbmm2 (version 1.16.99), which is a PacBio wrapper for the MiniMap2 aligner (version 2.26). A small fraction (mean 0.086%) of reads, having *<*70% gap-compressed identity with the reference genome (GRCh38), remained unmapped. We hypothesised that reads that have little identity with the reference may contain SVs and reanalyzed these unmapped reads to find local alignments.

Local alignments could directly correspond to inversions or translocations or be adjacent to insertions or deletions. We searched the human reference genome for two sets of *ε*-matches (Table 7) and found that a small number of alignments overlap a known SV. There were at least an order of magnitude more *ε*-matches that placed the query read to a region with a known *ε*-match. Due to alignment hotspots in repeats, we binned the alignments into 1000bp reference regions and found additionally between 21% and 54.8% of 1000bp bins that contained *ε*-matches but no known SVs.

**Table 7.** Results of finding local alignments for unmapped reads. We assume that unmapped reads remained unmapped due to larger rearrangements, which can be found by local alignment. Local alignments represent the structural variant directly or can be found for the rest of the long read. Known SV: The number of known structural variants that overlap a local alignment. Known SV region: The number of known structural variants in a region where at least one read was aligned. New SV: 1000bp regions that contain local alignments but no known structural variants.

| Sample | Unmapped count | $\varepsilon(50, 1)$ | | | $\varepsilon(100, 3)$ | | |
| --- | --- | --- | --- | --- | --- | --- | --- |
|  |  | Known SV | Known SV region | New SV (1000bp) | Known SV | Known SV region | New SV (1000bp) |
| HG00731 | 13680 | 470 | 6394 | 7763 (54.8%) | 223 | 3165 | 2823 (47.0%) |
| HG00732 | 22456 | 352 | 12631 | 7077 (35.9%) | 177 | 8343 | 2268 (21.3%) |
| NA19238 | 1088 | 192 | 2974 | 3224 (52.9%) | 102 | 1423 | 1101 (43.6%) |
| NA19239 | 4034 | 467 | 6290 | 4853 (43.5%) | 171 | 3165 | 1564 (33.0%) |

For this analysis, we used native DREAM-Stellar repeat masking with a 50% repetitiveness filter i.e for queries that had IBF hits in most reference regions, we searched in only the top 50% of the most variable (least repetitive) regions of the human genome.

To verify our predictions, we used Leaf [27], which is a tool for finding a preliminary list of likely structural variants. The example of an inverted translocation predicted on 2p from DREAM-Stellar alignments agrees with the variant detected by Leaf (Fig. 8).

**Fig. 8.**
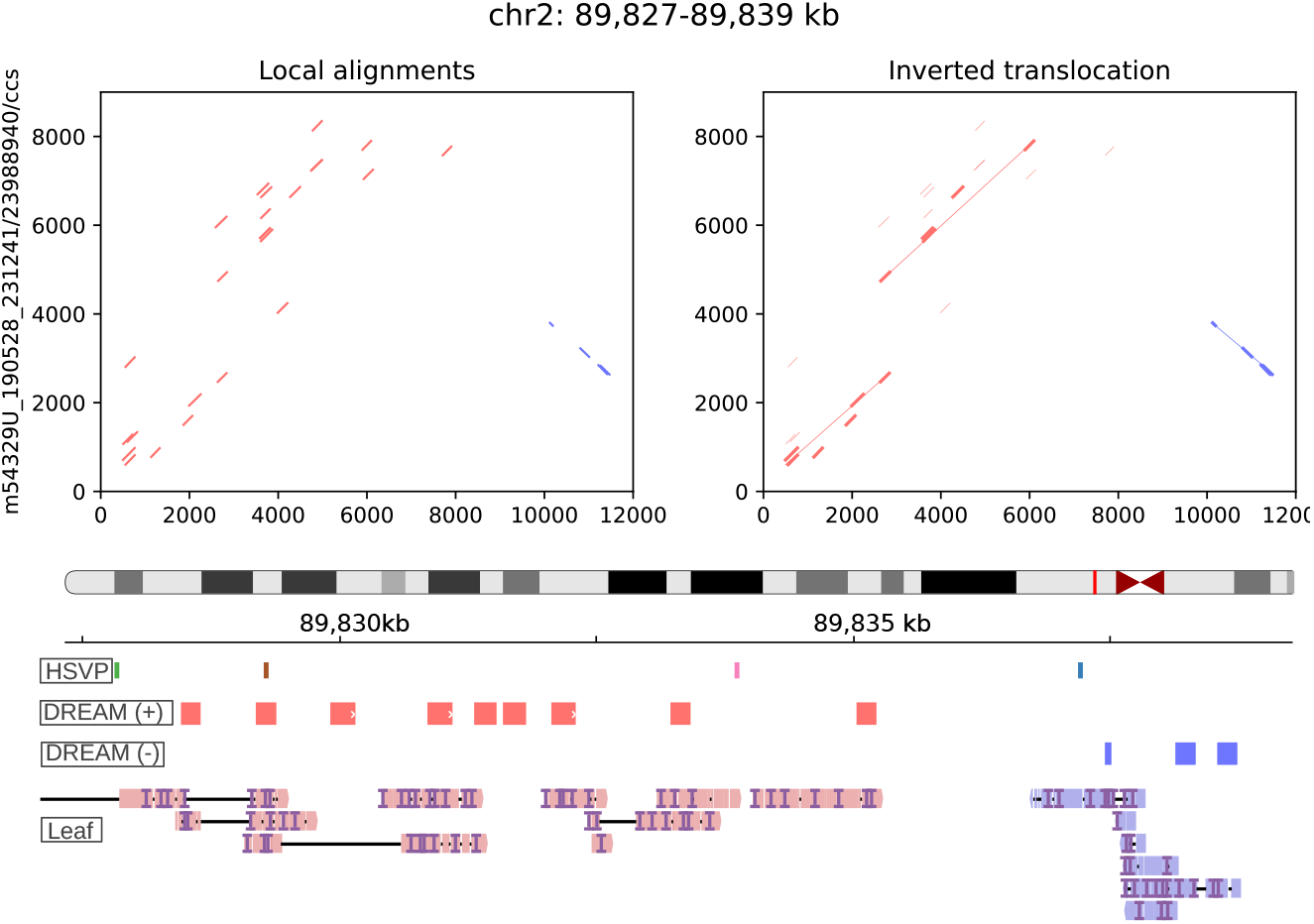
Above, the set of local alignments between chr2 and an unmapped PacBio HiFi read. The local alignments, split between the sense and missense strands, suggest an inverted translocation. The DREAM-Stellar alignments for the candidate structural variant are shown in the proximal portion of 2p. **HSVP**, variants from the Human Structural Variant Project; a deletion in HG00731 is shown in blue. **DREAM**, local alignments for the PacBio read on the sense (+) and missense (-) strands. **Leaf**, possible genomic arrangements for the read mapping on the sense (salmon) and missense (blue) strands.

## 4 Discussion

With DREAM-stellar we offer an efficient and accurate tool for approximate sequence search. More specifically, we can compare pairs of large sequences to find all conserved subsequences, i.e., the set of pairwise local alignments.

Seeding is a common first step in sequence search. Initially, an inexpensive, approximate method is applied to find a superset of potential alignment locations. In DREAM-Stellar, we apply two levels of increasingly specific k-mer-based prefiltering to narrow down the search space before alignment. In the DREAM approach, which we used for local alignment for the first time, we build an IBF over the reference and apply a k-mer filter. The benefit of adding the IBF prefilter to the existing Stellar aligner is two-fold. First, the IBF k-mer filter greatly reduces the search space before the SWIFT filtering in Stellar. Second, both sequences are split and distributed for parallel processing, which was not possible in Stellar.

The original single-threaded version of Stellar required almost 15h to find the ca. 5k *ε*(50, 3)-matches between the mouse and fly reference genomes. Through the use of the IBF k-mer filter in DREAM-Stellar, this search now only requires 1 min on 32 threads. This 900x speedup is not explained by the additional threads alone. DREAM-Stellar also compares favorably to a naively parallelized version of the alignment that does not apply prefiltering. A distributed Stellar search on 32 threads requires more than 1h and uses 101GB of main memory compared to the 13.5GB memory peak in DREAM-Stellar. Further, the benchmarks showed that only with efficient prefiltering could we find the set of *ε*-matches between large mammalian genomes.

The Stellar SWIFT filter stores the exact position of each query k-mer. The IBF filter indexes the reference k-mers instead and only stores whether a k-mer appears in a longer segment of the reference. Both filters apply a shared k-mer count threshold based on the *ε*-match definition. Storing the exact locations for the lossless SWIFT filter is more computationally costly and the SWIFT filter is more specific because the threshold is applied within a short *l* length region of the sequence. In the IBF filtering, the count of shared k-mers is obtained across a reference bin that is ca. 1000 times longer. To offset the spurious matches in the IBF filtering, we apply a lossy heuristic threshold for high error rates. However, the benchmarks showed that through a careful parameter selection, the number of missed matches in the heuristic k-mer filtering is small.

In k-mer filtering, we aim to identify local alignments based on k-mer matches. K-mer choice must strike a balance between long k-mers that can uniquely identify regions of interest (specificity) and short k-mers that are not destroyed by edits (sensitivity). The number of distinct k-mers and the expected specificity decrease exponentially with a linear decrease in k-mer length. If *k* is small, then all distinct *k*-mers are expected to appear in a large random sequence. In practice, we found that gapped *k*-mers, in particular symmetrical shapes, work well. The sensitivity of the gapped 18-mer 111010101101010111 is bounded by the sensitivity of its longest ungapped sub-mer 111 from above and the ungapped *k*-mer of the same weight 111111111111 (*w* = 12) from below. The actual sensitivity for a given gapped shape is not known; we use a pessimistic estimate, which ensures that the chosen k-mer weight and threshold lead to an accurate search. Our model estimates the expected FPR and FNR based on independent and uniformly random nucleotides. Of course, biological sequences are not random strings, as can be seen, e.g., by the existence of nullomers [28] and the more than 60k conserved CTCF binding sites [29].

Sequence search is hard because queries from repetitive regions have thousands of matches, which is detrimental to runtime. All the tools in our benchmark have a subroutine to handle repeats. BLAST masks locally repetitive subsequences using Dust. Similarly, Stellar has a filter for discarding repeats based on the local sequence context. LASTZ has a global approach for discarding words that appear often. LAST employs adaptive seeding where the seed length is increased until the number of seed hits is under some threshold. Additionally, LAST applies a threshold for the number of initial matches per query position, which we found has a significant effect on the number of matches and the runtime. In DREAM-Stellar, we order IBF bins based on the variability within a bin and avoid searching in repetitive bins if a query matches in more than half of all bins. If we were not able to reduce the search space by at least 50% we search in only the most variable bins among the query hits. We found that in GRCh38, highly repetitive bins correlate with centromeres, the Y chromosome, and extra-chromosomal parts of the human reference genome. In GRCh38, these repetitive regions are not confidently assembled, although there are efforts to do so in later versions [30]. We bias the search against repetitive regions to avoid excessive runtime and redundant matches.

An initial comparison using the native repeat masking in BLAST and Stellar showed that BLAST finds only ca 10% of Stellar *ε*-matches. The reason is that repeats are treated differently between tools and can make up the majority of the results. In the present version, we applied Dust repeat masking before searching with any of the aligners. Even then, for some benchmarks, there was a large difference in the number of alignments between the tools. DREAM-Stellar was as fast or faster than the other tools in the comparison. However, DREAM-Stellar specializes in relatively low error rates and yields less diverged alignments than BLAST, LAST, and LASTZ.

DREAM-Stellar, as the original Stellar, scores alignments based on a normalized edit distance and applies a minimum length criterion. Knowing the epsilon match definition and the sequence sizes, we can find the maximum E-value for an epsilon match. For the benchmarks in real genomes, we chose an edit distance and minimum length cutoffs that would lead to only alignments with a low E-value (*ε*(50, 3) in mouse vs fly *E <* 6·5 10^−7^ and *ε*(100, 6) in human vs mouse *E <* 7.5·10^−29^). Still, there exist alignments that have a low E-value, which are not *ε*-matches due to length or edit distance. When searching between two mammalian genomes that are relatively closely related, we found that the stringent *ε*-match criteria yielded a biologically relevant set of alignments. When searching for highly diverged matches, affine gap costs can be better suited than an edit distance-based score. Affine gap costs have implications for the alignment [31] and k-mer based prefiltering, where more shared k-mers are retained if edits appear in clusters (indels) compared to single edits distributed uniformly. Extending DREAM-Stellar to allow affine gap costs could extend the range of feasible error rates for local alignments.

## 5 Conclusion

We assessed our tool DREAM-Stellar on various input data from least (pair of simulated random sequences) to most similar (clipped alignment of reads to reference). We model the FPR and FNR statistics to deduce the suitable parameter values for the k-mer filter. DREAM-Stellar, with the programmatically deduced parameters, is fast and accurate in finding local alignments of a minimum length and maximum edit distance. The set of results contains many alignments that other heuristic tools cannot find with standard parameters, or, when the parameters are relaxed, are drowned in millions of other, not so significant results.

## Availability and requirements

**Project name:** DREAM-Stellar

**Project home page:** https://github.com/seqan/dream-stellar

**Operating systems:** Linux and MacOS

**Programming language:** C++ 20

**Other requirements:** Recent g++ or CLANG compiler and CMake to compile from source.

**License:** BSD 3

**Any restrictions to use by non-academics:** license needed

## List of Abbreviations

DREAM: Dynamic seaRchablE pArallel coMpressed index
IBF: Interleaved Bloom Filter
OOM: Out of memory
FPR: False positive rate
FNR: False negative rate
SINES: Short interspersed nuclear elements
LINES: Long interspersed nuclear elements

## Declarations

### Ethics approval and consent to participate

Not applicable.

### Consent for publication

Not applicable.

### Availability of data and material

Code used to produce the results and figures is available at https://github.com/eaasna/dream-stellar-benchmark. DREAM-Stellar v2.1.0 was used for the benchmarks in this manuscript and can be found on bioconda. PacBio HiFi sequencing data from Phase II of the Human Structural Variation Project for samples HG00731, HG00732, NA19238, and NA19239 are available at https://www.internationalgenome.org/data-portal/data-collection/hgsvc2.

### Authors’ contributions

EA led the development of the software, conceptualized the thresholding, conducted the benchmarks, and wrote the manuscript. ME assisted in refactoring the legacy codebase during the early stages of the project. SGG assisted in debugging during development and contributed the design of the shopping cart queue. KR conceived the project and revised the manuscript.

## Acknowledgements

We are sincerely grateful to Enrico Seiler for his valuable advice on software development and alignment-free search. This software makes use of the model of minimiser statistics developed by Enrico Seiler for the search tool Raptor (https://github.com/seqan/raptor).

## Appendix A

### Repeat masking

The Dust repeat masker scans a sequence for windows of 64 bases that contain few distinct 3-mers. Windows with distinct 3-mer count below a threshold are replaced with the unknown N base. We randomly sample from the DNA alphabet to replace these stretches of Ns. The number of distinct k-mers in the fly genome increases after repeat masking (RM Fig. 3). We used this hard masking approach in the comparison with other tools to make the results comparable.

### Threshold FNR estimate

For estimating the false negative rate of our approach, we introduce a simple model that proved to be sufficient for our purposes.

#### Problem definition

Given a sequence *S* of length *n* and another sequence, obtained by introducing *e* substitutions, insertions or deletions uniformly randomly across the original sequence, find the probability of the two sequences sharing at least *t* k-mers.

We find the expected FNR for the following parameters:

- *n* sequence length (epsilon min length)
- *e* edit distance (epsilon max errors)
- *t* threshold for minimum number of shared k-mers
- *k* k-mer length

Let A be the event that two sequences share at least *t* k-mers. The total number of k-mers in the sequence is *N* = *n* − *k* + 1. If A occurs, then the errors destroyed no more than (*N* − *t*) k-mers. If A does not occur, then in the complementary event *B* errors destroy at least (*N* − *t* + 1) k-mers.

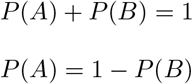

The number of ways in which errors can be distributed across the sequence is:

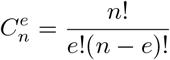

The probability of event B is the fraction of all error configurations that destroy at least *d* = (*N* − *t* + 1) shared k-mers. Let *f* (*k, t, e, l*) be a function that returns the number of error configurations that destroy at least *d* k-mers. We precalculate *f* (*k, t, e, l*) where

1. *k* ∈ [7, 23]
2. *t* ∈ [1, 25]
3. *e* ∈ [0, 15]
4. *l* ∈ [0, 150]

to find the expected number of false negatives. The false negatives are stored in a 3D dynamic programming matrix that contains a table for each possible threshold value.

Indexes are one-based and *S*[1…*n*] is the maximal subsequence of sequence *S* of length *n*.

**Base case: t = 1**

First, we consider the case where *t* = 1 and find all error configurations that destroy (*N* − 1 + 1) = *N* shared k-mers, i.e, all cases where sequences share no k-mers after mutation. Notice that the condition of no shared k-mers means that the leftmost error should be positioned in the range *S*[1…*k*]. Thus, the number of error configurations destroying all k-mers can be found as:

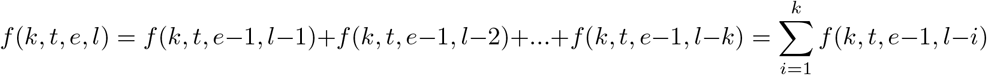

Fig. A 1 visualizes this sum in a table of varying error count and sequence length. Fig. A 2 highlights cases where all or none of the error configurations destroy all shared k-mers. Fig. A 3 shows how the calculation of *f* can be simplified.

**Fig. A1.**
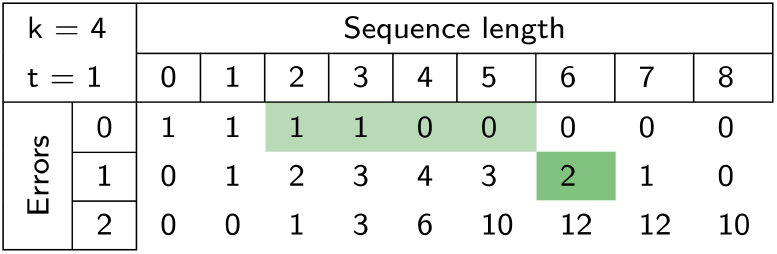
The number of error configurations that destroy all shared k-mers for *k* = 4 and *t* = 1. The value in the dark green cell can be found as the sum of the highlighted subvector in the previous row.

**Fig. A2.**
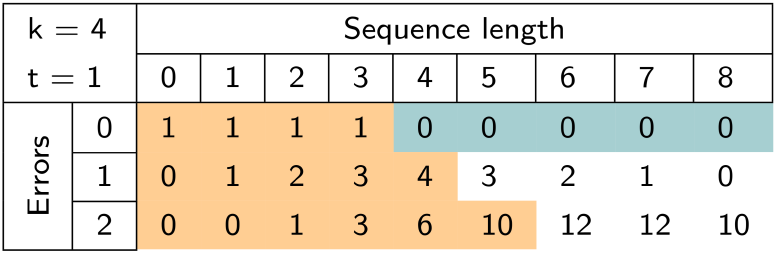
The values highlighted in blue are initialized to 0 if there are 0 errors in a sequence of length ≥ *k* meaning that no error configuration can destroy shared k-mers. The values in orange are 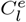 for the case when all possible error configurations destroy all shared k-mers.

**Fig. A3.**
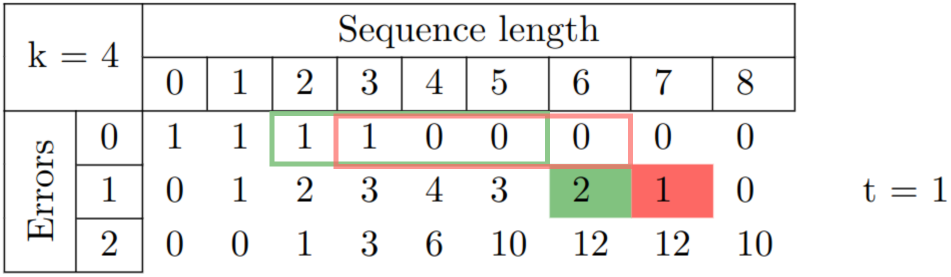
Instead of finding the sum of the subvector the red value can instead be calculated as *f* (4, 1, 1, 7) = *f* (4, 1, 1, 6) + *f* (4, 1, 0, 6) − *f* (4, 1, 0, 2).

#### General case

Next, we extend the base case by noting that if *t >* 1 then *f* (*k, t, e, l*) is the number of error configurations that destroy fewer than *N* shared k-mers, and the leftmost error can be at position [1…(*k* + *t*)]. In this case, the number of error configurations can be found as:

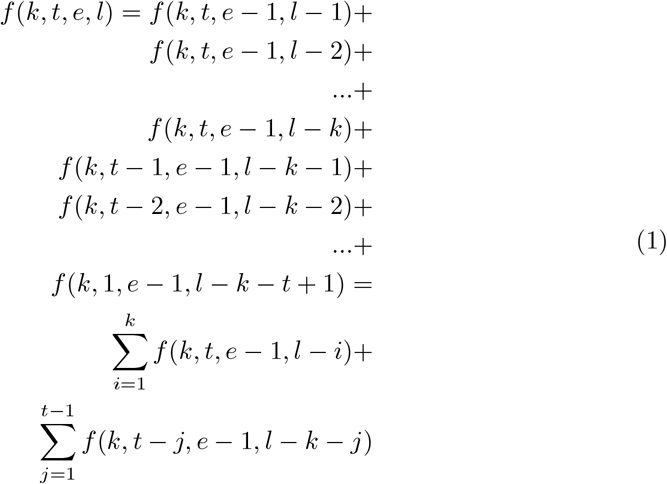

**Fig. A4.**
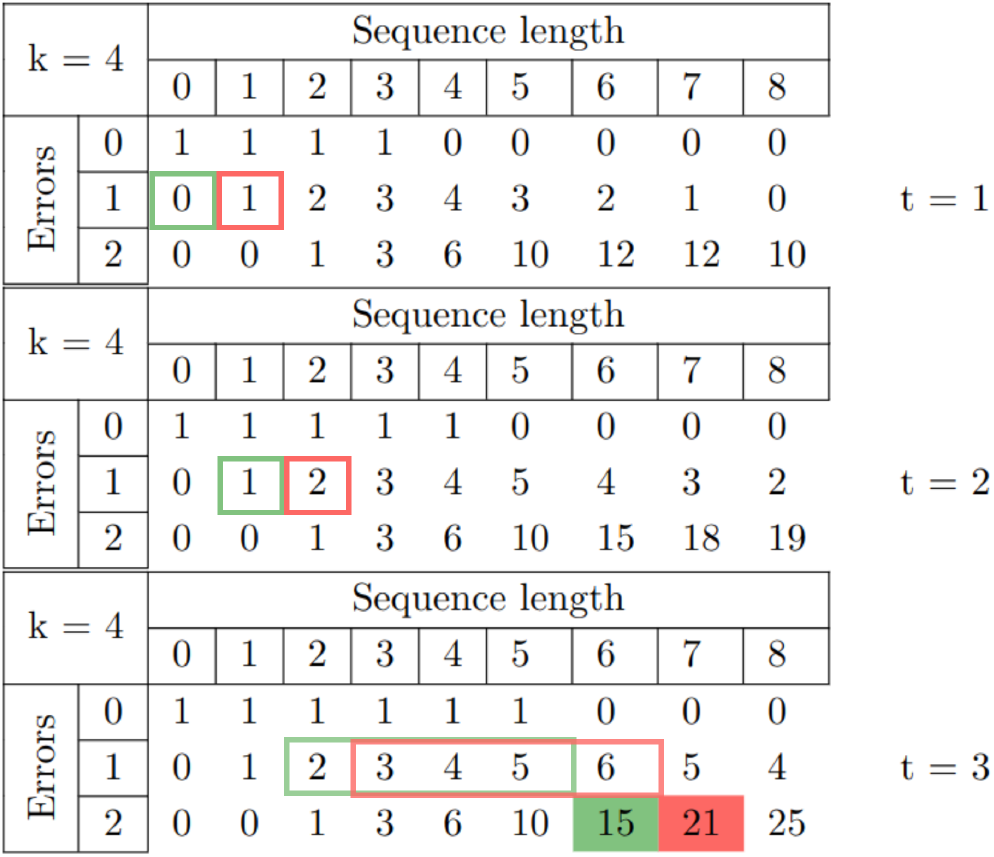
Analogously to Fig. A 3 the number of error configurations *f* (*k* = 4, *t* = 3, *e* = 2, *l* = 6) in green is the sum of a subvector in the previous row. Additionally, cases where the leftmost error appears after one *f* (*k* = 4, *t* = 2, *e* = 1, *l* = 1) or two *f* (*k* = 4, *t* = 1, *e* = 1, *l* = 0) matching k-mers have to be added.

Fig. A 4 visualises this sum in the 3D dynamic programming matrix. For an *ε*(6, 2)-match that has minimum length and maximum edit distance, there are 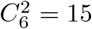 ways in which 2 errors can be distributed across 6 positions. A sequence of length 6 contains

*N* = 6 − 4 + 1 = 5 k-mers of length 4. All 15 error configurations destroy more than *N* − *t* = 5 − 3 = 2 k-mers (green cell Fig. A 4) and the expected false negative probability for *t* = 3 is 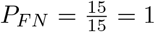.

### Threshold FPR estimate

In the alignment-free prefiltering strategy, we gauge the likelihood of reference and query segments sharing an *ε*-match based on the number of shared k-mers. False positive matches have to be validated, which increases the search runtime.

#### Hash collisions

A Bloom filter is a probabilistic data structure with a false positive rate that depends on the size of the filter, the number of hash functions, and the number of inserted elements. Due to hash collisions, there is a probability that a k-mer membership query returns true when the k-mer was not inserted into the bitvector. Let *P*_*IBF*_ be the false positive probability of a k-mer query. The higher bound for *P*_*IBF*_ is set at IBF construction based on the maximum number of elements in any reference bin.

#### Spurious matches

To estimate the suitability of different k-mer sizes, we model a sequence of length *L* as the set of k-mers. A sequence of length *L* has a total of *L* − *k* + 1 k-mers. Let *X* be the random variable representing the number of occurrences of some canonical k-mer when sampling a uniform distribution *L* − *k* + 1 times. There are |*σ*| ^*k*^ distinct k-mers in an alphabet of size |*σ*| ; half of these can be chosen as the canonical k-mer. Thus, *X* has a binomial distribution and a success probability of 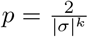 (right panel Fig. 3). Strictly, only bases and not k-mers are sampled independently in sequence simulation, but the dependence on adjacent overlapping k-mers is negligible when *k* ≪ *n*.

The expected number of occurrences for a canonical k-mer in a random DNA sequence is 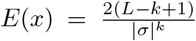. In a repetitive sequences k-mers are drawn from a smaller set and given the effective sequence size coefficient *s*_*e*_ the expected number of occurrences is 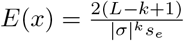.

We estimate the probability of a k-mer match in the case of independent and random sequences, considering the size of the reference database as *P*_*spur*_ = *min*(1, *E*(*x*)).

*P*_*IBF*_ and *P*_*spur*_ are not mutually exclusive, and the probability of a false positive k-mer query is thus *P*_*k*_ = *P*_*IBF*_ + *P*_*spur*_ −*P*_*IBF*_ *P*_*spur*_.

Partially overlapping patterns of minimum length are considered as seeds in the search. The number of k-mers in a match of minimum length *l* is *#k* = *l − k* + 1. The probability of a pattern exceeding the shared k-mer threshold when no eps-match exists is

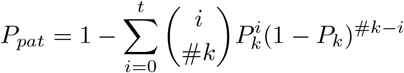

where *i* is the number of positive k-mer queries for a pattern. Given the false positive probability of a pattern *P*_*pat*_ and the number of patterns per query segment *#pat*_*seg*_, the probability that no patterns of a segment match is

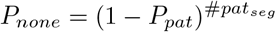

and the false positive probability of a segment is *P*_*seg*_ = 1 − *P*_*none*_.

A query segment is searched in each reference bin with a pattern match. False positives should be limited because they are detrimental to runtime. A careful parameter choice limits the segment false positive rate *P*_*seg*_ and thus the runtime.

### Parameter deduction

For a given reference database and some minimum length *l*_*min*_ and maximum normalized edit distance *e*_*max*_ we minimize

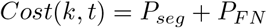

where *k* ∈ 7..23 and *t* ∈ 1..20 to find the suitable k-mer size and threshold for the IBF prefiltering.

Henceforth, the k-mer size *k* is fixed. Next, we find the k-mer hash values for the chosen *k* and index them into an IBF. The IBF is specific for a k-mer shape and minimiser scheme. The implications of shaped k-mers and nontrivial minimiser schemes are clarified below. For some dataset, the IBF can be built once and then reused for searches where *l* ≤ *l*_*min*_ and 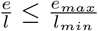. That means that an index built for *ε*(100, 10) can be used to find *ε*(50, 4).

Given the k-mer shape that was deduced for the IBF and some chosen *ε*-match definition, we find the threshold for the search.

Parameter deduction:

- Input:
  – Reference sequences
  – Distribution granularity (IBF bin count)
  – Query sequences
  – Eps-match definition (min length, max errors)
- Decision boundaries:
  – FPR upper bound *γ*
  – Threshold lower bound *t*_*lim*_
  – FNR upper bound *ν*
- Output:
  – Search kind ∈ {*lemma, gapped, heuristic, stellar*}
  – K-mer shape
  – Threshold

When searching for long alignments with a low error rate, we apply the k-mer counting lemma threshold. The k-mer counting lemma states that given two sequences that share a local alignment of length *l* that differ by *e* edit operations, the two sequences will share at least *t*_*lemma*_ = *l* − *k*(*e* + 1) − 1 k-mers. However, if for some FPR upper limit *γ* and threshold lower limit *t*_*lim*_

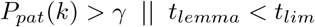

we use a heuristic k-mer threshold instead of the k-mer lemma threshold. For the heuristic threshold, we find the maximum *t*_*h*_ so that *P*_*FN*_ (*k, t*_*h*_) *< ν* for some FNR upper limit *ν*.

#### Gapped k-mers

Gapped k-mers are k-mers with some do-not-care positions. We represent k-mers using binary shape strings, e.g., 11111 is an ungapped 5-mer and 11011 is a k-mer of length 5 with a gap at the middle position.

There are |*σ*| ^4^ distinct 11011 k-mers in an alphabet of size |*σ*|, where 4 is the weight, i.e, the number of 1’s in the shape. All k-mer shapes of the same weight have an equal number of distinct k-mers for a given alphabet. For this reason, we equate the FPR statistics of a gapped k-mer with the FPR of an ungapped shape of the same weight.

In the worst case, a gapped shape has the same sensitivity as an ungapped shape of the same weight. There exists a configuration of errors that destroys the same number of gapped and ungapped k-mers. However, in practice, gapped shapes are more sensitive, especially if adjacent k-mers share few bases.

For prefiltering with gapped shapes, we apply a lossless k-mer lemma threshold if the error rate is relatively low. For a gapped k-mer shape of length *s* and weight *g*, the k-mer counting lemma states that two sequences that share an *ε*(*l, e*)-match share at least *t*_*lemma*_ = *l* −*s* + 1 −*e* · *g* k-mers. For higher error rates where the k-mer lemma threshold approaches 0, we apply a heuristic threshold and assume, based on the independence of adjacent k-mers, that more than the k-mer lemma lower bound number of gapped k-mers will remain after mutation.

#### Minimisers

The same sequence information is encoded in adjacent k-mers multiple times. For efficiency, it is beneficial to subsample the k-mer set to find a representative set of k-mers. DREAM-Stellar supports (*w, k*)-minimiser schemes where only the k-mer with the smallest hash value in a window of width *w* is added to the representative set. To find the k-mer hash value, we map k-mers into a base-four 64-bit space and XOR them with a random seed to avoid a skewed sampling towards A-heavy k-mers.

The number of (*w, k*)-minimisers of a sequence of fixed length is sequence-dependent and varies between some upper and lower bounds. To adjust a k-mer lemma threshold, we use the minimiser model from Raptor [16]. In case a heuristic threshold was chosen, we count the number of minimisers in patterns of the query to find the fraction of the upper bound and multiply the chosen threshold by the fraction.

**Fig. A5.**
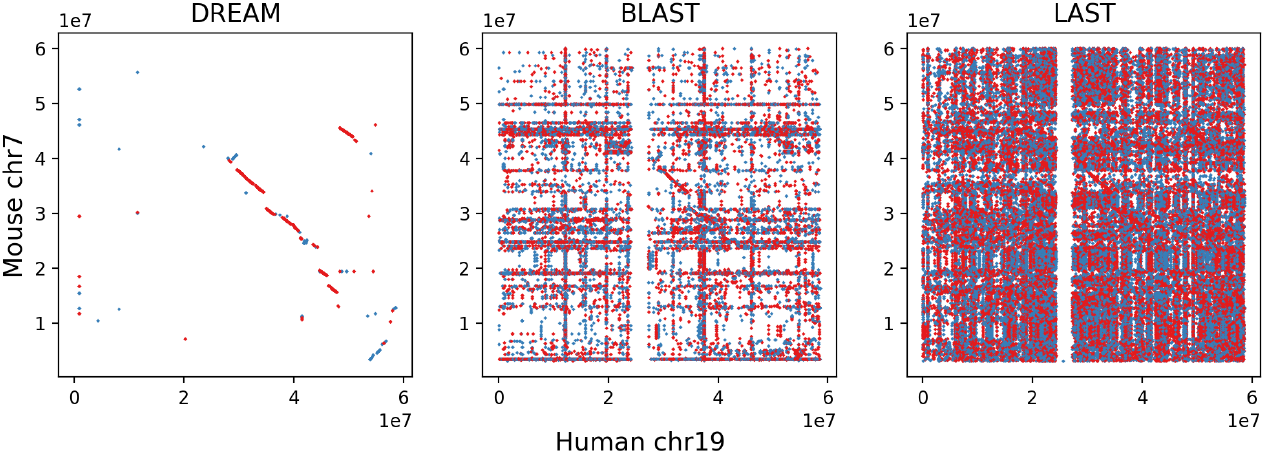
The set of DREAM-Stellar *ε*(100, 6)-matches illustrates the large contiguous blocks of homology between human chromosome 19 and mouse chromosome 7. Just as in chromosome 1, the mouse chromosome is inverted.

## Further results

### Reference genome indexing

#### Structural variants

The large blocks of homology between q19 in human and p7 in mouse are long established [32] are are also illustrated by the set of *ε*-matches shown in Fig. A 5.

